# ETV4 and AP1 transcription factors form multivalent interactions with three sites on the MED25 activator-interacting domain

**DOI:** 10.1101/126458

**Authors:** Simon L. Currie, Jedediah J. Doane, Kathryn S. Evans, Niraja Bhachech, Bethany J. Madison, Desmond K. W. Lau, Lawrence P. McIntosh, Jack J. Skalicky, Kathleen A. Clark, Barbara J. Graves

**Affiliations:** Department of Oncological Sciences, University of Utah School of Medicine, Salt Lake City UT, 84112-5500, USA.; Huntsman Cancer Institute, University of Utah, Salt Lake City UT, 84112-5500 USA.; Departments of Biochemistry and Molecular Biology, Department of Chemistry, and Michael Smith Laboratories, University of British Columbia, Vancouver BC, V6T 1Z3, Canada.; Department of Biochemistry, University of Utah School of Medicine, Salt Lake City UT, 84112-5650, USA.; Howard Hughes Medical Institute, Chevy Chase MD, 20815-6789, USA.

## Abstract

The recruitment of transcriptional cofactors by sequence-specific transcription factors challenges the basis of high affinity and selective interactions. Extending previous studies that the N-terminal activation domain (AD) of ETV5 interacts with Mediator subunit 25 (MED25), we establish that similar, aromatic-rich motifs located both in the AD and in the DNA-binding domain (DBD) of the related ETS factor ETV4 interact with MED25. These ETV4 regions bind MED25 independently, display distinct kinetics, and combine to contribute to a high-affinity interaction of full-length ETV4 with MED25. Within the ETS family, high-affinity interactions with MED25 are specific for the ETV1/4/5 subfamily as other ETS factors display weaker or no detectable binding. The AD binds to a single site on MED25 and the DBD interacts with three MED25 sites, allowing for simultaneous binding of both domains in full-length ETV4. MED25 also stimulates the *in vitro* DNA binding activity of ETV4 by relieving autoinhibition. ETV1/4/5 factors are often overexpressed in prostate cancer and genome-wide studies in a prostate cancer cell line indicate that ETV4 and MED25 occupy enhancers that are enriched for ETS-binding sequences and are both functionally important for the transcription of genes regulated by these enhancers. AP1-binding sequences were observed in MED25-occupied regions and JUN/FOS also contact MED25; FOS strongly binds to the same MED25 site as ETV4 AD and JUN interacts with the other two MED25 sites. In summary, we describe features of the multivalent ETV4- and AP1-MED25 interactions, thereby implicating these factors in the recruitment of MED25 to transcriptional control elements.

## Introduction

The activation domains (ADs) of sequence specific DNA-binding transcription factors interact with general transcription factors, coactivators, and chromatin remodelers, in order to regulate the location and activity of RNA polymerase II (Pol II) [1]. These interactions are important for the foundation of transcriptional programs that regulate development and establish cell-type identity [2]; as such, components of these interactions are commonly mutated in human disease [3, 4]. Acidic ADs, originally noted for an enrichment of negatively-charged and non-polar residues [5, 6], have an alternating pattern of negatively-charged/nonpolar and bulky hydrophobic/aromatic residues. Although usually disordered in isolation, ADs often become more helical when interacting with cofactors [7-10]. These sequence and structural characteristics are presumably the foundation of the ability of a single AD to interact with multiple partners as a flexible hydrophobic/aromatic interface that can be presented differently to diverse proteins [11-20]. However, higher affinity for a particular factor, and thus specificity, can be accomplished through the use of multiple ADs [21-24].

ETV1, ETV4, and ETV5 form a subgroup within the ETS family of transcription factors, sharing high sequence conservation both within and beyond the DNA-binding domain (DBD). This subgroup is aberrantly overexpressed in a subset of prostate cancers [25-27], and promotes PI3-kinase and RAS signaling pathways resulting in an aggressive and metastatic disease phenotype [28, 29]. Previously it was demonstrated that the N-terminal AD of ETV5 binds to the <u>ac</u>tivator interacting <u>d</u>omain (ACID) of Mediator subunit 25 (MED25) [7, 30]. However, ∼ 10% of prostate cancers frequently harbor truncations of ETV1, ETV4, or ETV5 that lack this AD due to chromosomal rearrangements [31, 32]. This suggests that the AD is dispensable for the function of these factors in prostate cancer. Therefore, we hypothesized that ETV1/4/5 subfamily factors could use an additional MED25-binding site, outside of the N-terminal AD, to interact with MED25. Such a domain, if it functioned in the absence of the AD, might explain the retained transcriptional activity of the oncogenic ETV1/ETV4/ETV5 truncations.

The Mediator complex is a critical transcriptional coactivator that serves as a primary conduit for transmitting regulatory signals from specific transcription factors to Pol II [33]. The 26 subunits of Mediator (not including the CDK8 kinase module) form distinct modules termed the head, middle, and tail. A reconstituted complex comprised of 15 subunits from the head and middle modules represents the minimal functional, or “core”, complex required for the general coactivator function of Mediator [34-36]. In contrast, the presence of, and requirement for, other subunits of Mediator, is more variable and gene-specific [37-47]. The simplest model to explain gene-specificity is that non-core Mediator subunits are required only for the transcription of the genes to which they are directly recruited via interactions with distinct sequence-specific transcription factors [33, 38, 40, 48, 49]. For example, in addition to ETV5 the transcription factors ATF6a, HNF4a, RARa, SOX9, and the viral protein VP16 recruit the variable subunit MED25 to their respective target genes [30, 43, 50-53]. The SOX9-MED25 interaction is implicated in chondrogenesis because reduced expression of either component results in similar palatal malformations in zebrafish [53]. Therefore, Mediator subunits can have gene-specific and cell-specific functions.

Here we investigate the biochemical basis and functional implications of ETV4-MED25 interactions. High-affinity interaction with MED25 was specific to the ETV1/4/5 subfamily, whereas, other ETS factors bound to MED25 with lower affinity, or not at all. Further investigation of the ETV4-MED25 interaction revealed that ETV4 has not one, but two domains that contacted MED25. Both regions, the N-terminal activation domain (AD) and the DNA-binding domain (DBD), bound the <u>ac</u>tivator <u>i</u>nteracting <u>d</u>omain (ACID) of MED25 via a similar motif (ΩxxxΩΦ or ΦΩxxxΩ, where Ω is an aromatic residue, Φ is a hydrophobic residue, and x is any residue). MED25 activated the DNA binding of ETV4 by relieving a previously described autoinhibition mechanism. Full-length ETV4, bearing both regions, had higher affinity for MED25 than either domain alone. Furthermore, the kinetics of association and dissociation by each domain differed, suggesting a complex binding reaction when both are present. NMR spectroscopy, mutational studies, and protein-docking modeling provided evidence that the AD and DBD bound to the same site on MED25 ACID. However, the DBD also interacted with two additional, distinct sites on MED25, such that simultaneous occupancy was possible in spite of overlapping contact surfaces. We provided *in vivo* evidence for the importance of this interaction as MED25 and ETV4 occupancies were highly overlapping genome-wide, and there was a significant overlap in genes whose expression was affected by depletion of either factor. MED25-occupied regions were enriched for ETS and AP1 binding sequences, and JUN/FOS heterodimers also contacted MED25 through a similar mechanism as ETV4. Therefore, we propose that both ETV4 and AP1 transcription factors use multivalent interactions to recruit MED25 to gene regulatory regions and promote the stable assembly of transcriptional machinery.

## Results

### High-affinity interaction with MED25 is specific to the ETV1/4/5 subfamily of ETS factors

The interactions between MED25 ACID (residues 391-553) and several ETS transcription factors were measured by biolayer interferometry in which one species is attached to a substrate and solution binding of an analyte is monitored (Fig. 1a,b). Testing single concentrations of these full-length ETS factors, we observed a range of interaction strengths with MED25. ETV1 and ETV4 at 50 nM were sufficient for interaction with MED25, whereas, tenfold more ETS1 and SPDEF (500 nM) were required. No interaction with MED25 was detected with 500 nM of EHF, ERG, or ETV6. We measured the relative strength of ETS factors binding to MED25 more accurately by determining the equilibrium dissociation constants (*K_D_*) from kinetic rate constants (*k_a_* and *k_d_*) (Fig. 1c, Fig. S1a,b, and Table S1). Interestingly, for all ETS factors the interaction data better fit a model assuming two ETS proteins binding to MED25, rather than a one-to-one interaction (Fig. S1c). As other experimental approaches, discussed below, also support a multivalent ETV4-MED25 interaction, we report values calculated using the two-to-one model. ETV4, with K_D_ values of 7 ± 3 and 28 ± 7 nM for the two interactions with MED25, and ETV1 (16 ± 5 and 21 ± 2 nM) bound to MED25 with 20- to 50-fold higher affinity compared to SPDEF (320 ± 90 and 5,000 ± 2,000 nM) (Fig. 1c). Therefore, we conclude that high-affinity interaction with MED25 is specific to the ETV1/4/5 subfamily of ETS transcription factors.

**Fig. 1.**
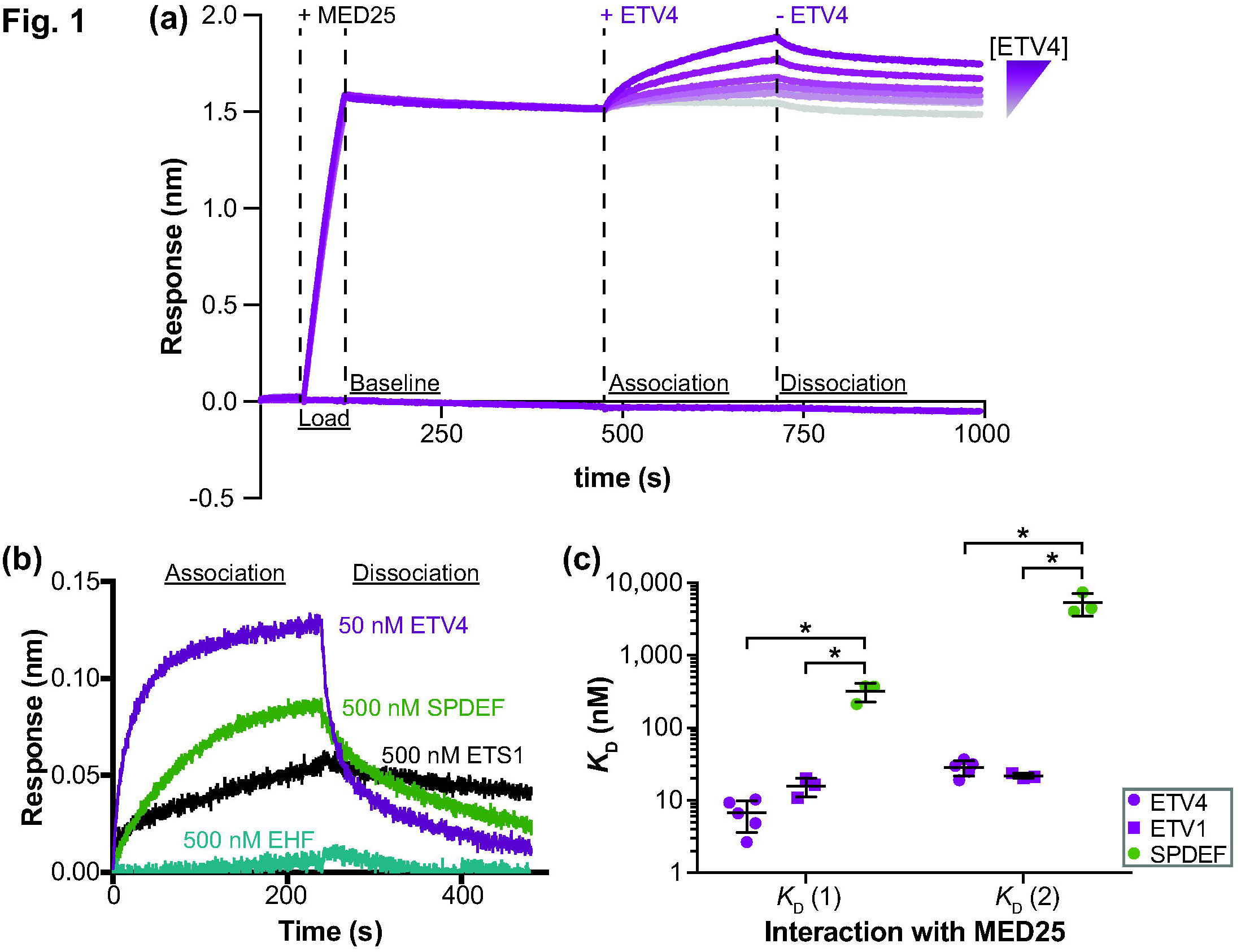
High-affinity interaction with MED25 is specific to the ETV1/4/5 subfamily of ETS factors. (a) Representative ETV4-MED25 sensorgrams using biolayer interferometry. Streptavidin sensors were loaded with the biotinylated ACID domain of MED25^391-553^ (+ MED25), followed by the addition (+ ETV4), then removal (-ETV4) of six different concentrations of ETV4 in order to measure the association and dissociation, respectively, between ETV4 and MED25 ACID. The bottom purple line is a control sensor with no MED25 loaded and the highest concentration of ETV4 added during the association step. (b) Representative sensorgrams of binding assays with a single concentration of ETV4 (50 nM), SPDEF (500 nM), ETS1 (500 nM), and EHF (500 nM). Only the association and dissociation phases of the sensorgram are shown. ERG (500 nM) and ETV6 (500 nM) were also tested but showed no detectable association; each ETS factor was tested twice. (c) Equilibrium dissociation constants (*K*_D_) for the interaction between MED25 ACID and SPDEF, ETV1, and ETV4. Two *K*_D_ values, denoted *K*_D_ (1) and *K*_D_ (2) with *K*_D_ (1) being the higher-affinity interaction, are reported as these interactions better fit a 2:1 (ETS:MED25) binding model (Fig. S1c). *K*_D_ values were determined using a group fit with six different concentrations of each ETS factor (Fig.1a and Fig. S1a,b). Circles and squares represent the *K*_D_ determined from a single, six-concentration experiment. Horizontal lines represent the mean and standard deviation for three to five replicate experiments. *K*_D_, *k_a_*, and *k*_d_ values for these interactions are summarized in Table S1. “*” Indicates *p* < 0.05 in a Mann-Whitney *U* test.

### Two distinct regions of ETV4 bind to MED25

We next sought to investigate the molecular basis of selectivity for high-affinity interaction between the ETV1/4/5 subfamily and MED25 ACID. Using ETV4 as a model for this subfamily, we interrogated the interaction between different fragments of ETV4 and MED25 with biolayer interferometry (Fig. 2a). The N-terminal AD, ETV4^43-84^, bound to MED25 in a one-to-one manner with a KD of 700 ± 100 nM (Fig. 2b,d and Table S1). This value is comparable to previous measurements of the interaction between the conserved AD of ETV5 and MED25 by fluorescence polarization (580 ± 20 nM) and by isothermal calorimetry (540 ± 40 nM) (Fig. S2) [7, 30]. As the AD bound MED25 with an approximately hundred-fold weaker affinity than full-length ETV4, we surmised that additional regions within ETV4 also contribute to the interaction with MED25. Indeed, ETV4^165-484^, which lacks the N-terminal AD, also bound to MED25 and used a two-to-one interaction mode reminiscent of full-length ETV4 (350 ± 80 and 2,200 ± 500 nM) (Fig. 2c,d). Testing of additional ETV4 fragments revealed that the minimal AD, ETV4^43-84^, and a broader N-terminal fragment, ETV4^1-164^, were equivalent in binding to MED25 (Fig. 2d). Likewise, ETV4^337-436^, which corresponds to the ETS domain and an additional α-helix H4 that is specific to the ETV1/4/5 subfamily, interacted with MED25 with similar affinity to that of ETV4^165-484^. Therefore, we conclude that the N-terminal AD and the C-terminal DBD contribute to the high-affinity binding of full-length ETV4 with MED25.

**Fig. 2.**
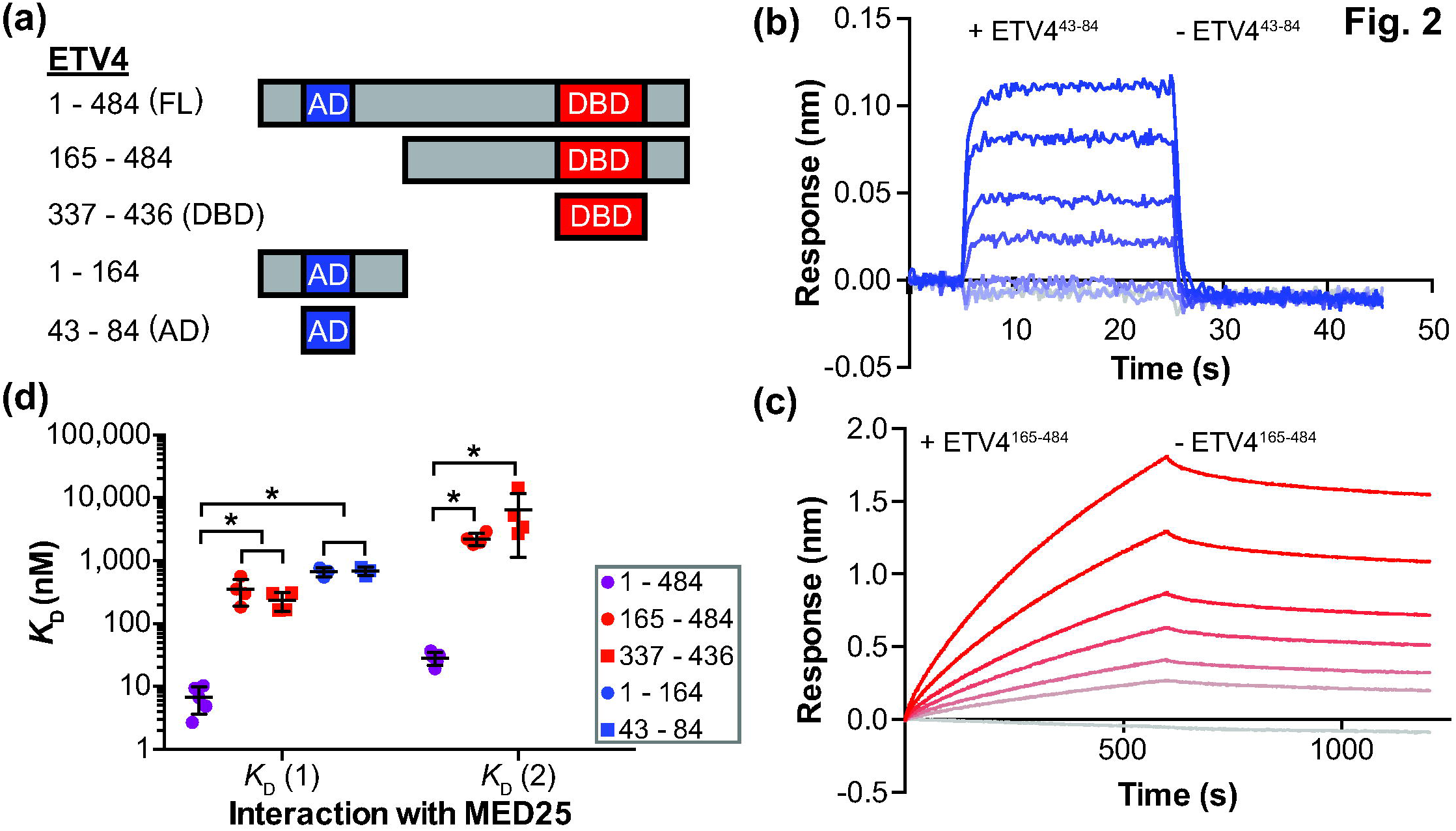
Two regions of ETV4 can independently bind to MED25. (a) Schematic of ETV4 truncations used for binding studies with MED25 ACID. AD, DBD, and FL are abbreviations for activation domain, DNA-binding domain, and full length, respectively. (b) Representative ETV4^43-84^-MED25 sensorgrams with the association and dissociation phases shown. 30 uM ETV4^43-84^ was used for the top sensorgram with serial 1.5-fold dilutions for the next five sensorgrams, and the bottom gray sensorgram corresponds to a control with no ETV4^43-84^. (c) Representative ETV4^165-484^-MED25 sensorgrams displayed as in (b). 3 uM ETV4^165-484^ was used for the top sensorgram. The difference in x- and y-axis scales between (b) and (c) reflects the different kinetic constants for these interactions (Table S1). (d) *K*_D_ values for the interactions between fragments of ETV4 and MED25 ACID. Filled circles and squares represent the *K*_D_ value from a single experiment with six different concentrations of ETV4, and the horizontal lines represent the mean and standard deviation for three to five replicate experiments. “*” Indicates *p* < 0.05. Note that there are two **K*_D_* values for ETV4^165-484^ and ETV4^337-436^ as these sensorgrams were better fit by a 2:1 (ETV4:MED25) model. In contrast, there is a single KD value for ETV4^1-164^ and ETV4^43-84^ as these sensorgrams were better fit by a 1:1 model. ETV4^1-484^ data are included from Fig. 1c for reference.

Interestingly, the kinetics of the ETV4 AD and DBD interactions with MED25 were noticeably different. The AD-MED25 interaction had relatively high association and dissociation rate constants (*k_a_* and *k*_d_, respectively), reflecting the faster association and dissociation for this interaction (Fig. 2b and Table S1). In contrast, the DBD-MED25 interaction had relatively low k_a_ and k_d_ values indicating slower association and dissociation (Fig. 2c and Table S1). These data suggest that the high affinity interaction between ETV4 and MED25 could be due to combining the fast association mechanism and the slow dissociation mechanism used by the AD and DBD, respectively.

### Interaction between MED25 and ETV4 DBD: molecular interface and influence on DNA binding

Previous studies provided structural characterization of the interaction between the ETV5 AD and MED25. Notably, the predominantly disordered AD becomes more helical in the MED25-bound state and phenylalanine and tryptophan residues in the AD are critical for this interaction [7, 30]. Based on the robust sequence conservation between the ADs of ETV1, ETV4, and ETV5 (Fig. S2), we surmised that the ETV4 AD would interact with MED25 in a conserved fashion. Indeed, the introduction of Phe54Ala, Phe60Ala, or Trp64Ala mutations into ETV4 significantly disrupted interaction with MED25 (Fig. S3a,b). Therefore, we focused on characterizing residues that are important for the newly discovered interaction between the DBD of ETV4 and MED25.

NMR spectroscopy was used to investigate the MED25-interface of the ETV4 DBD. We compared the ^15^N-HSQC spectra for ^15^N-labeled ETV4 DBD [54] with or without unlabeled MED25 ACID (Fig. S4a). MED25 most strongly perturbed the signals from residues (Glu425 and Ser429) within α-helix H4 of ETV4 (Fig. 3a and Fig. S4b,c). Additionally, signals from residues within H1, H3, and the β-sheet were perturbed. Based on the specificity of the ETV1/4/5 subfamily for high-affinity interactions with MED25, we focused on residues near the β-sheet and in H4. Surface-exposed residues on the β-strands, as well as residues in loops flanking the β-strands, are poorly conserved among all ETS factors (Fig. S2). Likewise, the sequence and secondary structure of H4 is not conserved in any ETS factors outside of the ETV1/4/5 subfamily (Fig. S2). To test the functional importance of individual residues we analyzed the influence of single site alanine substitutions on the ETV4-MED25 interaction. Mutations near the β-sheet (Thr363Ala, Lys370Ala, Ile372Ala, Glu373Ala) had a ∼ four- to ninefold disruption on MED25 binding (Fig. 3b and Fig. S3a). Mutation of Ser429 within H4 disrupted MED25 binding greater than tenfold. Ser429 occurs within a motif (<u>LF</u>SLA<u>F</u>) on H4 that is reminiscent of the N-terminal AD (<u>F</u>QET<u>WL</u>). This portion of the AD is critical for interaction with MED25 (Fig. S3a) and the conserved sequence in ETV5 becomes more helical when interacting with MED25 (Fig. S2) [7]. By analogy to the AD, we reasoned that the surface-exposed phenylalanines in H4 might also be important for interaction with MED25. Indeed, mutation of both Phe428 and Phe432 to alanine resulted in an approximately thirty-fold disruption of MED25 binding (Fig. S3a,c). These results suggest that a broad interface, including critical phenylalanines from α-helix H4 as well as residues from the β-strand of the ETS domain, contributes to the interaction of the DBD with MED25. Helix H4, by itself, resembles the AD suggesting that the AD and DBD of ETV4 bind MED25 through a similar motif (Fig. 3c). However, the contribution of additional residues outside of H4 suggests that the DBD uses a broader interface for interaction with MED25.

**Fig. 3.**
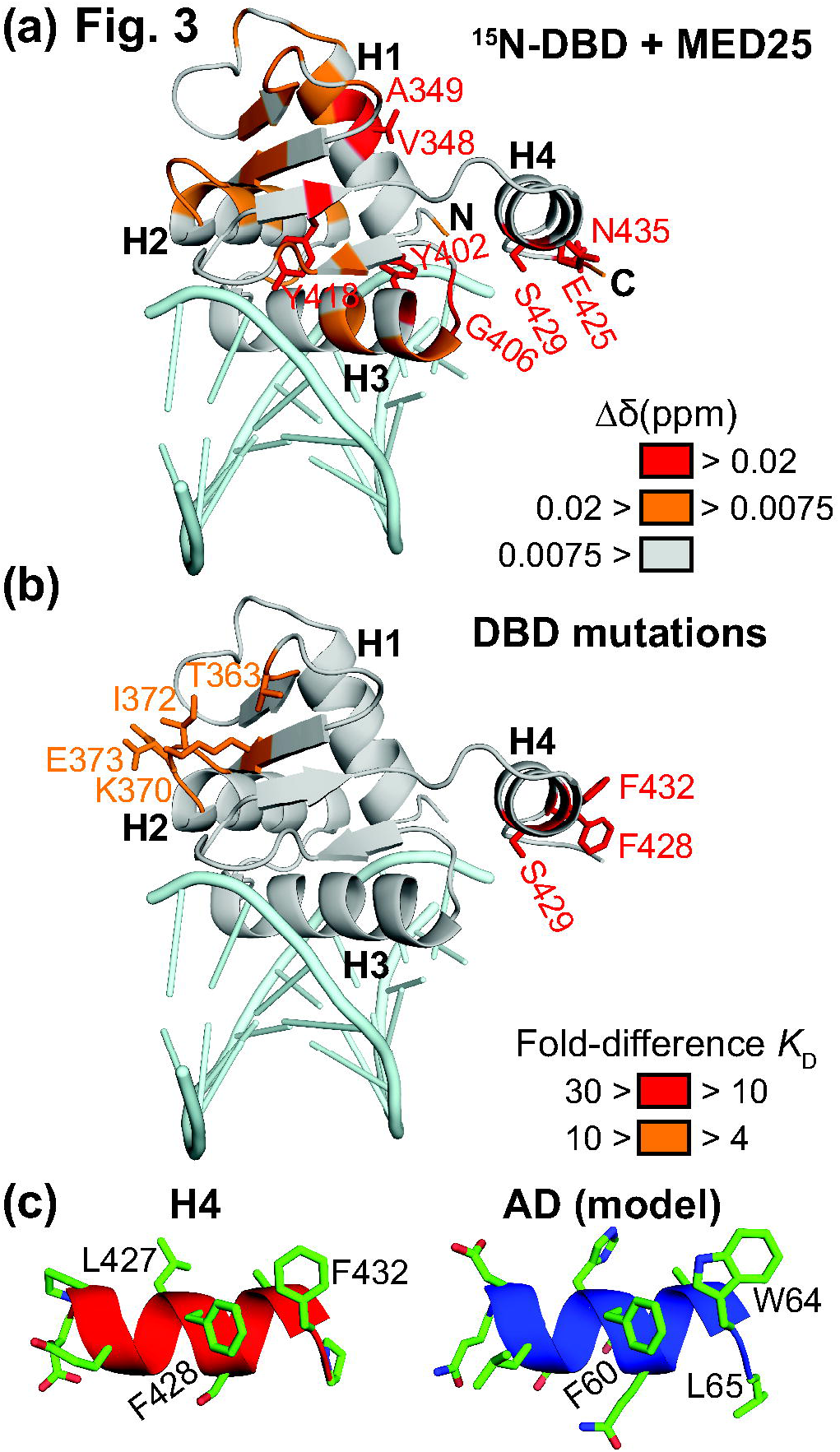
A similar motif in ETV4 AD and DBD is critical for binding to MED25. (a) Changes to the ^15^N-HSQC of ETV4 DBD upon addition of unlabeled MED25 ACID at a 1:1.2 molar ratio are colored onto the structure of ETV4 DBD bound to DNA (PDB: 4UUV) [56]. Amide chemical shift perturbations (Δδ = [(Δδ_H_)^2^ + (0.2Δδ_N_)^2^]^½^) that were greater than the mean are colored orange and those that were in the highest ten percent are colored in red. The peak for the amide of Glu425 is also colored red as this peak was broadened to baseline. DNA (light blue) is shown for illustrative purposes, although it was not included in the NMR experiment. See Fig. S4 for NMR spectra and quantification. (b) Fold-differences of *K*_D_ values for interaction with MED25 are mapped onto the DBD of ETV4. Indicated residues were mutated to alanine in full-length ETV4, and interaction with MED25 was compared to wild type ETV4. See Fig. S3 for example sensorgrams and quantification of ETV4 mutants. (c) Structure of H4, left, and model of AD, right, in cartoon representation with side chains shown in stick representation. Note that the AD is intrinsically disordered, but takes on partial helical character when interacting with MED25 [7]; this helical model for the AD was generated by swapping AD amino acids onto the structure of H4.

We previously demonstrated that α-helix H4 inhibits the DNA binding of the ETS domain of ETV4 [54]. Leu430 on the interior face of H4 is critical for this autoinhibition and interacts with conserved hydrophobic residues of the ETS domain. Hence, we questioned whether MED25 binding to the exterior face of H4 would affect DNA binding. Using an electrophoretic mobility shift assay (EMSA), we observed that the addition of MED25 to the equilibrium DNA binding reaction resulted in a slower migrating band on the EMSA gel, which we propose to be a MED25-ETV4-DNA ternary complex (Fig. 4a). As a control, equivalent amounts of MED25 did not interact with the EHF-DNA complex. The presence of MED25 increased the affinity (*K*_D_) of ETV4 for DNA approximately twofold (Fig. 4b). This matches the two-fold magnitude of DNA-binding autoinhibition that was previously attributed to H4 [54]. Mutation of the MED25-interaction site on H4 (Phe428Ala/Phe428Ala) or of the ETS domain-interaction site on H4 (Leu430Ala) both abrogated the activation of DNA binding by MED25 (Fig. 4b). We conclude that the interaction between MED25 and α-helix H4 activates the DNA binding of ETV4.

**Fig. 4.**
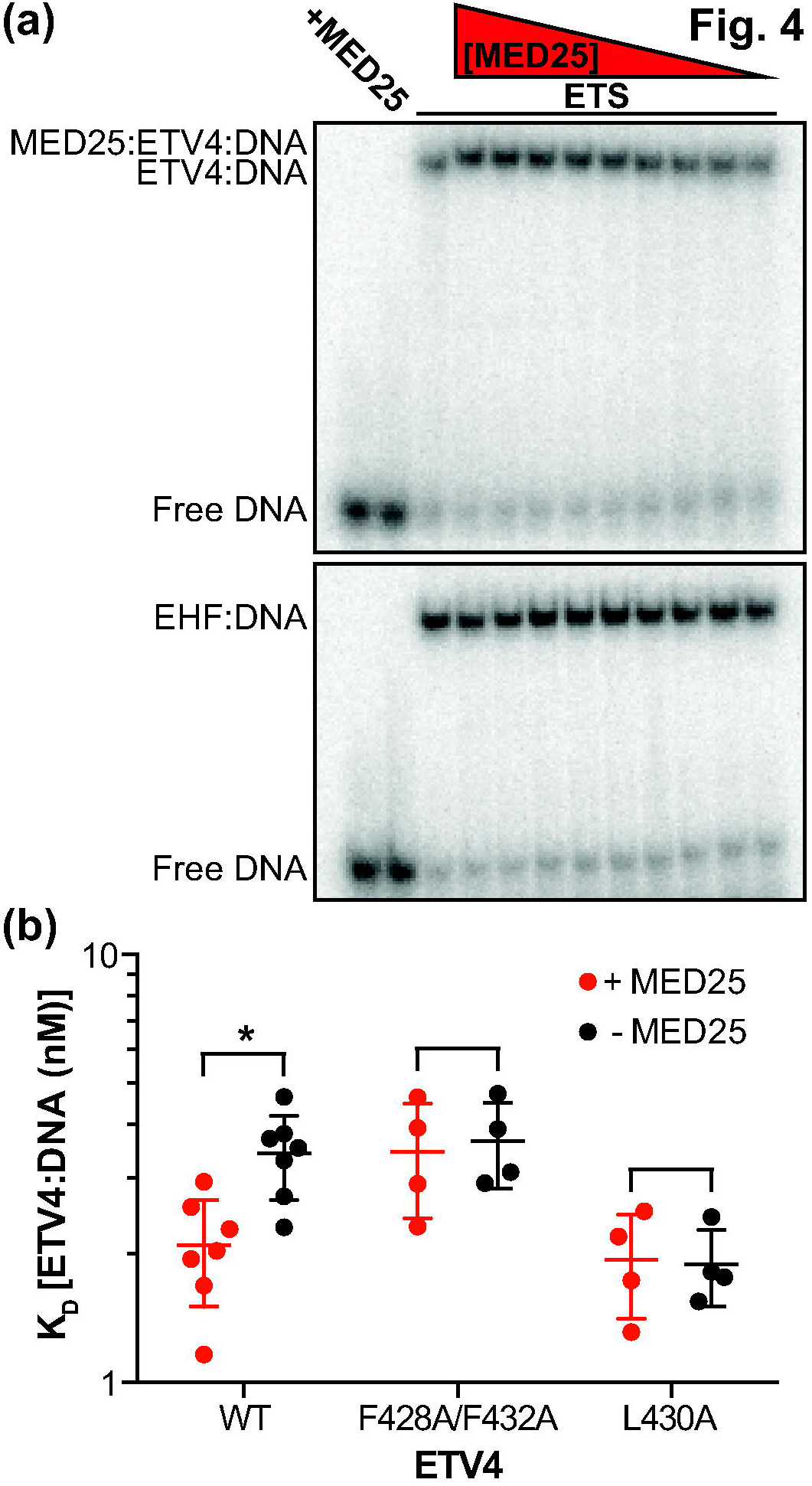
Interaction with MED25 relieves the DNA-binding autoinhibition from α-helix H4 of ETV4. (a) Electrophoretic mobility shift assay (EMSA) titrating MED25 ACID (500 – 1 nM) with a single concentration (50 nM) of ETV4 (top) or EHF (bottom). EMSA gels are representative examples of at least three replicates for each protein. (b) Using a single concentration of MED25 ACID (500 nM), ETV4 WT or indicated mutants were titrated (100 – 0.01 nM) to determine the K_D_ of the ETV4-DNA interaction in the presence or absence of MED25. Filled circles represent a single experiment and the horizontal lines represent the mean and the standard deviation. “*” Indicates *p* < 0.05. The *K*_D_ values for the ETV4-DNA interaction were (mean ± standard deviation): WT (+ MED25) = 2.1 ± 0.5 nM; WT (-MED25) = 3.4 ± 0.8 nM; Phe428Ala/Phe432Ala (+MED25) = 3.4 ± 1.0 nM; Phe428Ala/Phe432Ala (-MED25) = 3.7 ± 0.8 nM; Leu430Ala (+ MED25) = 1.9 ± 0.5 nM; Leu430Ala (-MED25) = 1.9 ± 0.4 nM.

### ETV4 AD and DBD interactions with MED25: single-site versus multisite binding

Having determined that the AD and DBD use similar motifs to interact with MED25, we next wanted to explore the AD and DBD binding sites on MED25. We utilized assigned ^15^N-HSQC spectra and the tertiary structure of the ACID domain of MED25 that were previously reported [9, 10]. The ACID domain is a seven-stranded β-barrel with three peripheral α-helices (Fig. 5a). ^15^N-labeled MED25 ACID was titrated with either the AD or the DBD of ETV4 (Fig. S5a,b). The addition of the AD resulted in robust and widespread changes in the ^15^N-HSQC spectra of MED25 ACID (Fig. S5c,e). Several amide ^1^N^H^-^15^N peaks displayed progressive chemical shift changes indicative of fast exchange behavior. We observed substantial line broadening, even at the lowest titration point, as has also been observed in MED25 ACID interactions with other activation domains [7, 9, 10]. Comparatively, the addition of ETV4 DBD resulted in more subtle changes in the spectra of MED25 ACID (Fig. S5d,f). Amide signals showed relatively smaller chemical shift changes and line broadening was more gradual as DBD was added at 0.2:1, 0.5:1, and 1.2:1 molar ratios. Although exhibiting different effects upon AD and DBD titration, we interpreted the changes in the MED25 ACID ^15^N-HSQC spectra as evidence for interaction with both of these ETV4 fragments.

**Fig. 5.**
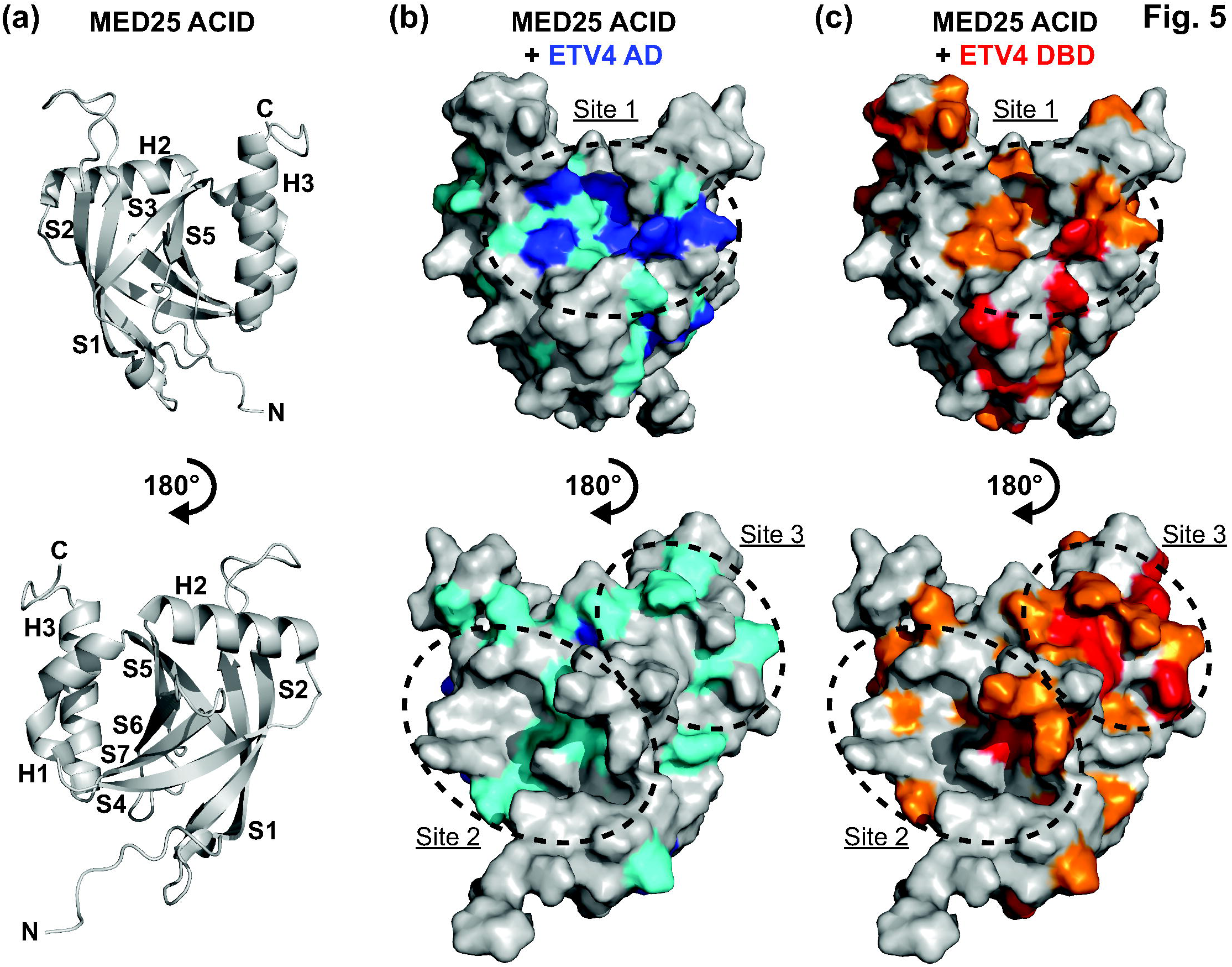
NMR spectroscopy suggests a single MED25 binding site for ETV4 AD and multiple binding sites for ETV4 DBD. (a) Cartoon representation of MED25 ACID structure (PDB: 2KY6) [9]. Bottom view is rotated 180° relative to the top view of MED25 ACID. α-helices and β-strands are abbreviated H and S, respectively, and numbered according to progression from N- to C-termini of the domain. (b) MED25 ACID oriented as in (a) but with surface representation and changes to the ^15^N-HSQC of MED25 ACID upon addition of unlabeled ETV4 AD indicated by color. Upon addition of 0.2 molar equivalents of ETV4 AD, MED25 amide relative peak intensities that were less than the mean are colored teal and those that were in the lowest ten percent are colored blue. (c) MED25 ACID as in (b) with changes to the ^15^N-HSQC upon addition of unlabeled ETV4 DBD indicated by color. Upon addition of 0.2 molar equivalents of ETV4 DBD, MED25 amide relative peak intensities that were less than the mean are colored orange and those that were in the lowest ten percent are colored in red. Sites of clustered changes upon titration of AD and/or DBD are indicated by dotted lines and arbitrarily named site one, two, or three. See Fig. S5 for ^15^N-HSQC spectra of MED25 ACID alone and with unlabeled ETV4 AD or ETV4 DBD.

We mapped the MED25 residues that exhibited amide peak intensity changes from both ETV4 titrations onto the structure of MED25 ACID (Fig. 5 and Fig. S5e,f). The AD most strongly perturbed residues that clustered in a single site formed by β-strands S3, S5, and α-helix H3, which we will term Site 1. The DBD influenced residues within Site 1 as well, but also at two different sites on additional faces of MED25 formed by S4, S7, and H1 (Site 2), and by S2, S4, and H2 (Site 3), respectively. These multiple sites all have concave grooves that are suitable for interaction with an α-helix, such as that formed by the AD or H4 of ETV4 (Fig. S6). Hydrophobic and uncharged polar residues form the floor of these grooves with positively charged residues lining the perimeter. While the AD appears to most strongly interact with Site 1 on MED25, we propose two possibilities to explain the absence of clustering for amide signal changes upon DBD titration to a single MED25 site. One possibility is that the DBD may alternately interact with multiple sites on different faces of MED25 ACID, thereby forming a “fuzzy” binding interface [8, 12]. Alternatively, the DBD may interact with a single site on MED25 and have significant through-molecule effects that influence other surfaces of MED25 without direct interaction. We next pursued further biochemical and structural characterization in an attempt to distinguish between these two models.

Mutagenesis was used to further investigate the binding of MED25 and ETV4. Using MED25 residues implicated from the NMR experiments as a starting point, we focused on those that had surface-exposed side chains. Residues from the concave groove floors, and surrounding positively charged residues, of all three MED25 ACID sites were mutated. Based on the importance of aromatic residues in the AD and DBD for binding to MED25 (Fig. 3a), most residues were mutated to glutamate as we surmised that a negative charge would successfully disrupt any π-cation or hydrophobic interactions. The strongest mutants for both the AD (Gln451Glu) and DBD (Lys422Glu, Arg509Glu, Arg538Glu, and Lys545Glu) were located at Site 1 on MED25, suggesting that this may be the preferred binding site for either of these ETV4 fragments in isolation (Fig. 6, Fig. S7, and Table S2). Interestingly, the mutations that disrupt AD binding cluster near the center of the groove in Site 1, whereas, the mutations that preferentially affect DBD binding are more spread out on Site 1. This suggests a broader surface at Site 1 is used for DBD binding, mirroring the broad surface on the DBD that is used for MED25 binding (Fig. 3a,b). Although some mutations in MED25 Site 2 and Site 3 also disrupted AD binding, all MED25 mutations at these sites (Gln430Glu, Arg466Glu, His474Glu, Tyr487Ser, Leu514Glu, Lys518Ala/Lys519Ala/Lys520Ala, Ile521Glu, and Met523Glu) more strongly, or only, affected DBD binding (Fig. 6 and Table S2). The spacing of the DBD-specific mutations at MED25 Sites 2 and 3 were also suggestive of a broad interface being important for interaction with the DBD at each of these sites.

**Fig. 6.**
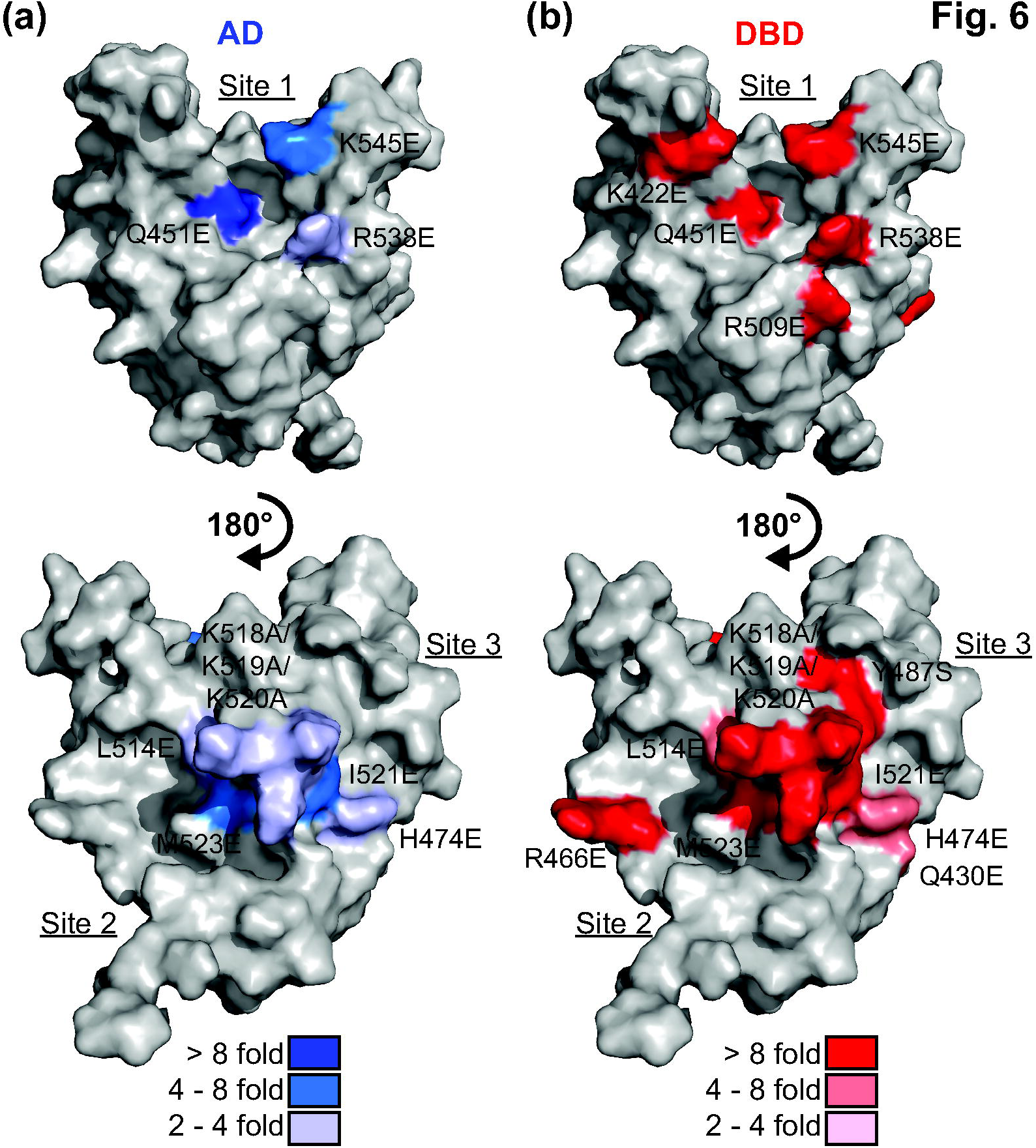
Broad surfaces on multiple MED25 sites contribute to DBD binding. MED25 ACID represented as in Fig. 5b and with point mutants that perturb ETV4 binding colored onto the structure. Blue scale (a) and red scale (b) indicate mutations that disrupted the interaction with ETV4 AD and DBD, respectively. Fold inhibition of binding was calculated by comparing to a wild type MED25 control with AD or DBD. See Table S2 for summary of all *K*_D_ values, and Fig. S7 for examples of sensorgrams with MED25 mutants.

Therefore, we interpret the importance of several surface-exposed residues at three distinct sites on MED25 ACID to support multiple MED25 binding sites for ETV4 DBD.

We used a protein-docking program, ZDOCK, to predict the structures of the AD-MED25 and DBD-MED25 bound complexes. ZDOCK uses complementary shape to analyze all possible binding modes between two proteins [55]. In addition to the structures for MED25 [9] and ETV4 DBD [56], we generated an α-helical model of the AD for input into the protein-docking program. Based on the mutational data for ETV4, we restrained the predictions to include Phe60 and Trp64 of the AD, and Phe428 and Phe432 of the DBD in the interface for each of these respective interactions. We also restrained the AD-MED25 prediction to include MED25 Gln451 because NMR clearly localized AD binding to Site 1 on MED25 and the Gln451Glu was the single strongest mutant for AD binding. No MED25 residues were required to be involved in the DBD-MED25 modeling due to the broad localization of both the NMR spectral perturbations of amides, and the mutational effects on DBD binding, to multiple faces of MED25. The top ten predictions for the AD-MED25 interaction are very similar with the AD clustering in MED25 Site 1 (Fig. 7a). The top ten DBD-MED25 predictions included the DBD binding to each of the three MED25 sites that were identified from NMR spectroscopy and mutational analysis, even though this information was not used to inform the predictions (Fig. 7b). In addition to α-helix H4, residues from H1 and the loops between β-strands on the DBD also contact MED25 in many of the predictions, thereby contributing to a broader interface with MED25. In summary, NMR spectroscopy, mutational analysis, and protein-docking predictions support multiple DBD binding sites on MED25 ACID.

**Fig. 7.**
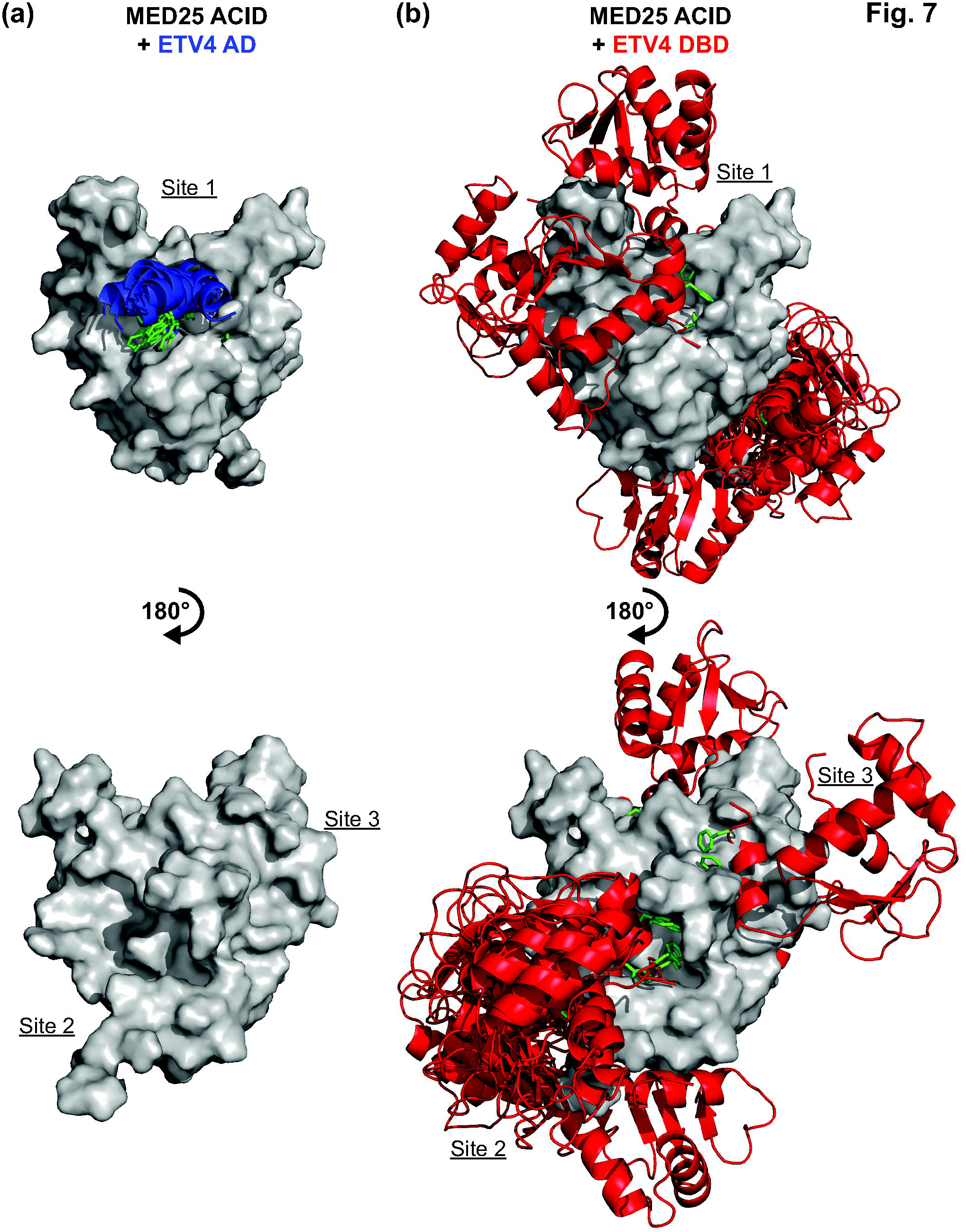
Modeling suggests multiple potential MED25 binding sites for ETV4 DBD. (a) The top ten predictions for the interaction between MED25 ACID and ETV4 AD using the ZDOCK protein-docking program [55]. MED25 ACID is represented as in Fig. 5b, ETV4 AD is displayed as a blue α-helix in cartoon representation with the side chains for Phe60 and Trp64 shown in green. AD residues Phe60 and Trp64 and MED25 residue Gln451 were selected as residues involved in the binding site for the prediction. (b) The top-ten predictions for the interaction between MED25 ACID and ETV4 DBD. The DBD was binding to MED25 Site 1 in two predictions, Site 2 in seven predictions, and Site 3 in one prediction. Modeling and representation were performed as in (a). The entire DBD is shown in cartoon format in red, and Phe428 and Phe432 side chains are displayed in green. DBD residues Phe428 and Phe432 were selected as residues involved in the binding site for the prediction.

### ETV4 and MED25 share transcriptional targets in prostate cancer cells

Given the strong interaction between MED25 and ETV4, as well as the established role of MED25 as a transcriptional co-regulator [30, 43, 50, 51], we hypothesized that MED25 and ETV4 would share transcriptional targets. To test this, we assayed ETV4 and MED25 DNA occupancy genome-wide in PC3 cells, a prostate cancer tumor cell line that overexpresses ETV4. Chromatin immunoprecipitation (ChIP) of a FLAG-tagged version of MED25 detected 1042 MED25 bound regions while the parallel ETV4 ChIP detected 847 ETV4 bound regions (Table S3). The vast majority of the enriched regions for each dataset mapped greater than 5,000 base pairs (bp) from defined transcriptional start sites, with almost half lying greater than 50,000 bp away, suggesting robust binding to distal enhancer elements (Fig. S8a,b). Intersection of the two datasets demonstrated a striking high degree of overlap; ∼75% of the ETV4 peaks were in the MED25 dataset (Fig. 8a,b and Table S3). We observed high enrichment of ETS binding motifs at the overlapping sites; the ETS binding site (CAGGAA) was the top overrepresented motif in the shared peaks and second top hit for all MED25 peaks after the AP1 binding sequence (Fig. 8c and Fig. S8c). We compared the frequency of the ETS binding motif between the shared peaks and a size-equivalent, randomly generated set of genomic sequences. Of the 611 ETV4-MED25 shared peaks, 223 had at least one C(C/A)GGAA sequence in their central 100 bp core; whereas, only 44 regions of a control dataset had the motif (Fig. 8c and Table S3). Furthermore, only two of the 44 regions in the control dataset had multiple occurrences of the motif, while 50 of the 223 shared peaks had two or more ETS motifs. Several targets were confirmed by qPCR to validate the ChIPseq data and verify that these represent overlapping binding regions (Fig. 8d and Fig. S8b).

**Fig. 8.**
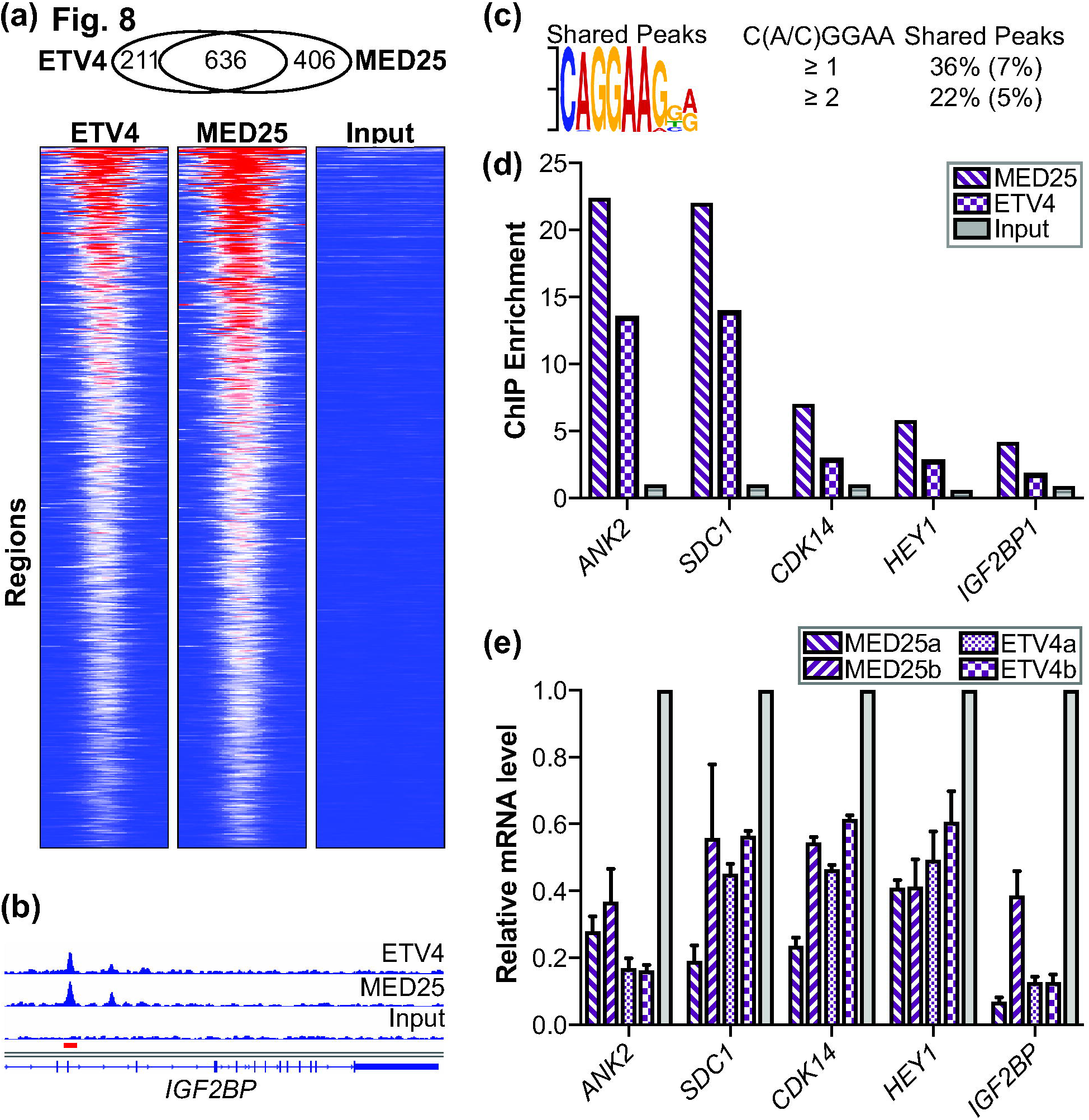
ETV4 and MED25 occupancy at regulatory regions genome-wide and effects on expression of associated genes. (a) Overlap of ETV4 and FLAG-MED25 bound regions from PC3 cells displayed as heatmaps of the RPKM from MED25, ETV4, and input control data across the set of shared regions. Heat map ranges from 0 (blue) to 72 (red) RPKM. (b) Graphical display of enriched reads for ETV4 and MED25 ChIP DNA and input DNA in IGF2BP intron (hg19 chr10:54,218,829-54,242,010). Red bar, region assayed by qPCR. (c) ETS binding motif is most over-represented sequence in shared regions as determined by MEME [81]. The MEME, expect-value, E, is 1 × 10^-21^. Percentages of peaks with match to motif are shown to right with comparison to randomly generated size-matched peak set in parentheses. (d) qPCR quantification of MED25-FLAG and ETV4 enrichment at putative regulatory elements for genes shown; input values displayed for comparison. Two to three independent biological replicates provided similar patterns, but different maximum levels of enrichment. A representative experiment is shown. (e) Relative expression values for indicated gene as determined by qPCR of total cDNA derived from ETV4 and MED25 knockdown PC3 cells. Two different shRNA constructs were used for both ETV4 and MED25 and denoted as a or b. Average values for biological triplicates are graphed with standard deviation. Relative expression of control knockdown cells is set at 1, and experimental sample values are graphed relative to control.

To test the functional relevance of this genomic occupancy, we sought to identify genes whose transcription was dependent on both MED25 and ETV4. We performed shRNA-mediated knockdown of endogenous MED25 and ETV4 in the PC3 cell line followed by RNAseq. Of the 153 and 57 genes down regulated by MED25 and ETV4 knockdown, respectively, 18% were in common (Table S5). Focusing on these common genes, 70% were associated with potential regulatory elements occupied by both ETV4 and MED25 (Fig. S8b). The occupancies of putative regulatory elements for five genes (*ANK2, CDK14, HEY1, IGF2BP1, and SDC1*) were confirmed by qPCR of DNA in the peak region (Fig. 8d and Fig. S8b). Consistent with RNAseq results, all five of these genes were down-regulated in PC3 cells expressing either MED25 or ETV4 shRNA construct as assessed by RT-qPCR (Fig. 8e). From these findings we propose that MED25 and ETV4 interact at genomic sites and work together to regulate the transcription of ETV4 target genes.

### JUN/FOS also binds to multiple sites on MED25

We were intrigued by the strong enrichment of AP1 binding sequences in our MED25 ChIP-seq dataset (Fig. S8c), as a MED25-AP1 interaction had not been previously reported. Using a published JUND ChIP-seq dataset from PC3 cells [57], we found a 29% overlap in MED25- and JUND-occupied regions (Table S3). Furthermore, JUN and FOS also had one and two of the MED25-interacting motifs (ΩXXXΩΦ), respectively, occurring within previously annotated activation domains (Fig. S9) [58, 59]. To investigate this potential interaction, we used full-length JUN and FOS to prepare JUN/FOS heterodimers, JUN homodimers, and FOS monomers. MED25 strongly bound FOS monomers (*K*_D_ = 9 ± 3 nM) (Fig. 9a, Fig. S10a, and Table S6). Although JUN dimers did not measurably interact with MED25 at a concentration of 500 nM, the presence of JUN in JUN/FOS heterodimers resulted in a slightly higher affinity for MED25 (*K*_D_ = 5 ± 3 nM) compared to FOS by itself (Fig. S10b). The apparent complex binding of JUN/FOS to MED25 was reminiscent of the AD and DBD of ETV4 combining to interact with MED25; the fast initial association of JUN/FOS was similar to FOS by itself, and the addition of JUN resulted in a second, slower association not observed for either JUN or FOS alone. This complex interaction behavior suggested that JUN/FOS also binds to multiple sites on MED25.

**Fig. 9.**
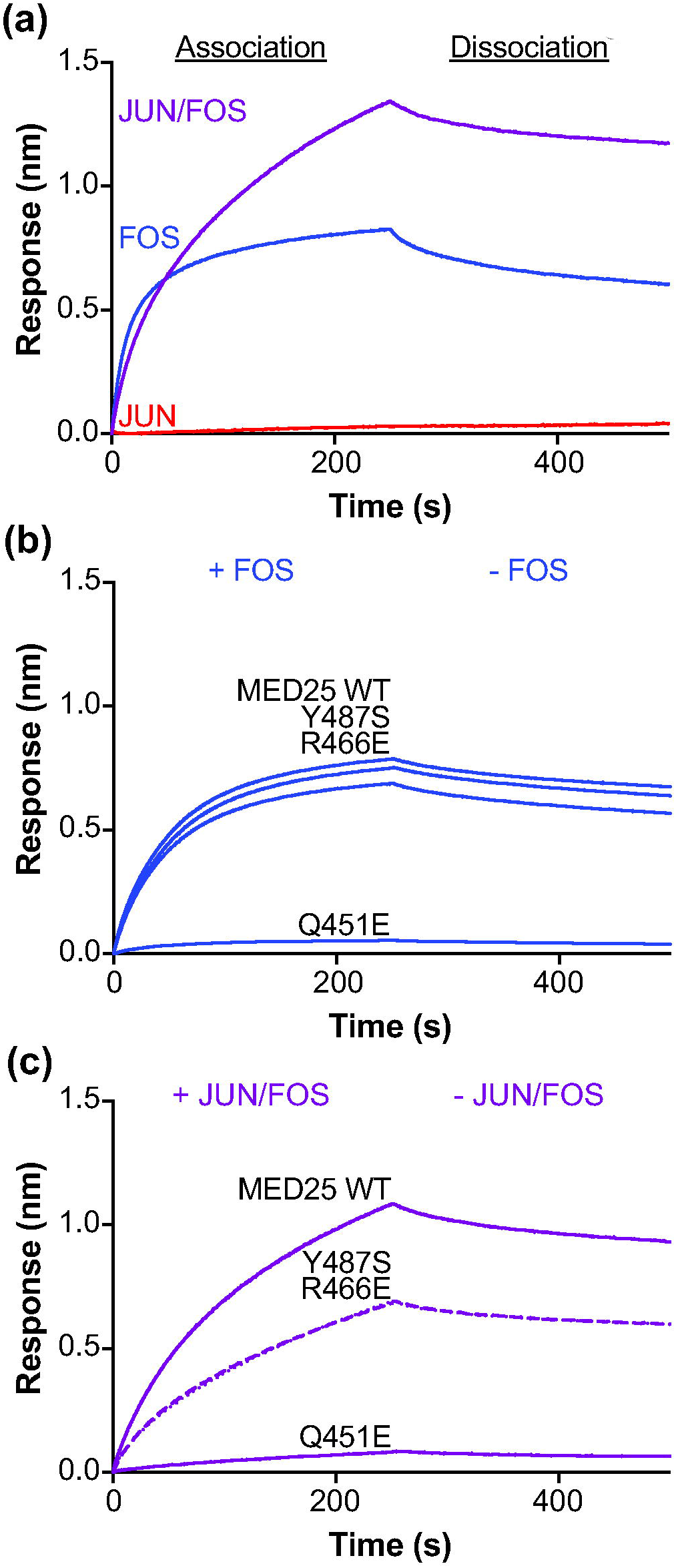
JUN/FOS heterodimers bind to multiple sites on MED25. (a) Representative sensorgrams of binding assays between MED25 WT and 500 nM JUN (red), 500 nM FOS (blue), or 250 nM JUN/FOS (purple). (b) Representative sensorgrams of binding assays between MED25 wildtype (WT) and point mutants as labeled, and 500 nM FOS. (c) Representative sensorgrams of binding assays between MED25 wildtype (WT) and point mutants as labeled, and 250 nM JUN/FOS. Note that sensorgrams corresponding to Tyr487Ser and Arg466Glu are shown as dotted and dashed lines, respectively, to aid in visualization because they are almost completely overlapping. See Fig. S10 and Table S6 for further quantification of JUN/FOS-MED25 and FOS-MED25 interactions.

We used MED25 mutants to further characterize JUN- and FOS-binding sites. Mutation of MED25 Site 1 (Gln451Glu) strongly disrupted the binding of both JUN/FOS and FOS alone (Fig. 9b,c and Table S6). In contrast, MED25 Site 2 (Arg466Glu) and Site 3 (Tyr487Ser) mutations had essentially no impact on FOS binding but disrupted JUN/FOS binding by two- to threefold. Based on these mutational data we propose that FOS within JUN/FOS heterodimers strongly binds to MED25 Site 1, and JUN weakly enhances the overall interaction by contacting Sites 2 and 3.

## Discussion

Here we report that both the DNA-binding domain (DBD) and the N-terminal activation domain (AD) of ETV4 bind to the activator interacting domain (ACID) of MED25. Both domains contribute to a higher-affinity interaction with MED25, as compared to ETS factors outside of the ETV1/4/5 subfamily. Three distinct sites on MED25 bind the DBD; whereas, only one site binds the AD, resulting in a multivalent interaction between MED25 and ETV4. MED25 activates the DNA binding of ETV4 by relieving autoinhibition, and we propose that the ETV4-MED25 interaction is important for the transcriptional output of ETV4 target genes. JUN/FOS heterodimers also interact with multiple sites on MED25, indicating that multivalent interactions may be a common mechanism of MED25 recruitment by diverse transcription factors.

### Model for ETV4- and JUN/FOS-MED25 interactions

The AD and DBD of ETV4 can each bind MED25 independently. Similar ΩXXXΩΦ or ΦΩxxxΩ motifs, where Ω represents an aromatic residue, Φ represents a hydrophobic residue and x represents any residue, are found in both domains and are critical for MED25 binding. However, this motif in the flexible AD is at the core of a single interaction surface; Phe60Ala and Trp64Ala mutations almost completely abrogate binding to MED25. By comparison, additional residues assist the interaction motif in the DBD and form a broader interface with MED25. This structural distinction correlates with the differences in kinetic rates for these interactions. We hypothesize that the flexible AD interface associates rapidly with MED25 but also dissociates quickly due to the lack of supporting interactions; whereas, the broader DBD interface orients more slowly during association and forms a more stable interaction with MED25 that dissociates more slowly.

Biochemical results from biolayer interferometry binding studies as well as structural insight from NMR spectroscopy and protein-docking predictions indicate that the DBD interacts with three distinct sites on MED25, and the AD interacts with only one of these sites. These three MED25 sites are relatively similar in overall appearance; hydrophobic and non-polar residues form the bottom of concave grooves with positively charged residues lining the perimeter. We speculate that the difference in single-site versus multisite binding for the AD- and DBD-MED25 interactions, respectively, reflects the different binding interfaces. For the AD, the exact spatial and chemical nature of the binding groove on MED25 Site 1 is critical for binding because this is the only point of contact. In contrast, the specifics of the MED25 binding grooves for binding to H4, at all three sites, are less critical for the overall interaction because other parts of the DBD also contact MED25 to form a broader interface. Based on the spacing of the sites on MED25 ACID, one DBD molecule could only interact with one MED25 site at a time. Yet, the presence of additional binding sites suggests that all three MED25 sites collectively contribute to the macroscopic DBD-MED25 interaction.

We propose a multi-step kinetic model for the ETV4-MED25 interaction (Fig. 10). An important assumption supported by our findings is that the AD and DBD cannot co-occupy the same MED25 site due to steric clash. The predominant pathway for complex formation would involve the AD of ETV4 first binding to Site 1 on MED25. The DBD of the bound ETV4 would then interact with MED25 at either Site 2 or Site 3 and could toggle between alternately binding to either of these two MED25 sites. The predominant pathway for dissociation will be limited by the slower release of the DBD. In this scheme, the AD and DBD of ETV4 and three MED25 ACID sites combine to form a high-affinity and multivalent interaction.

**Fig. 10.**
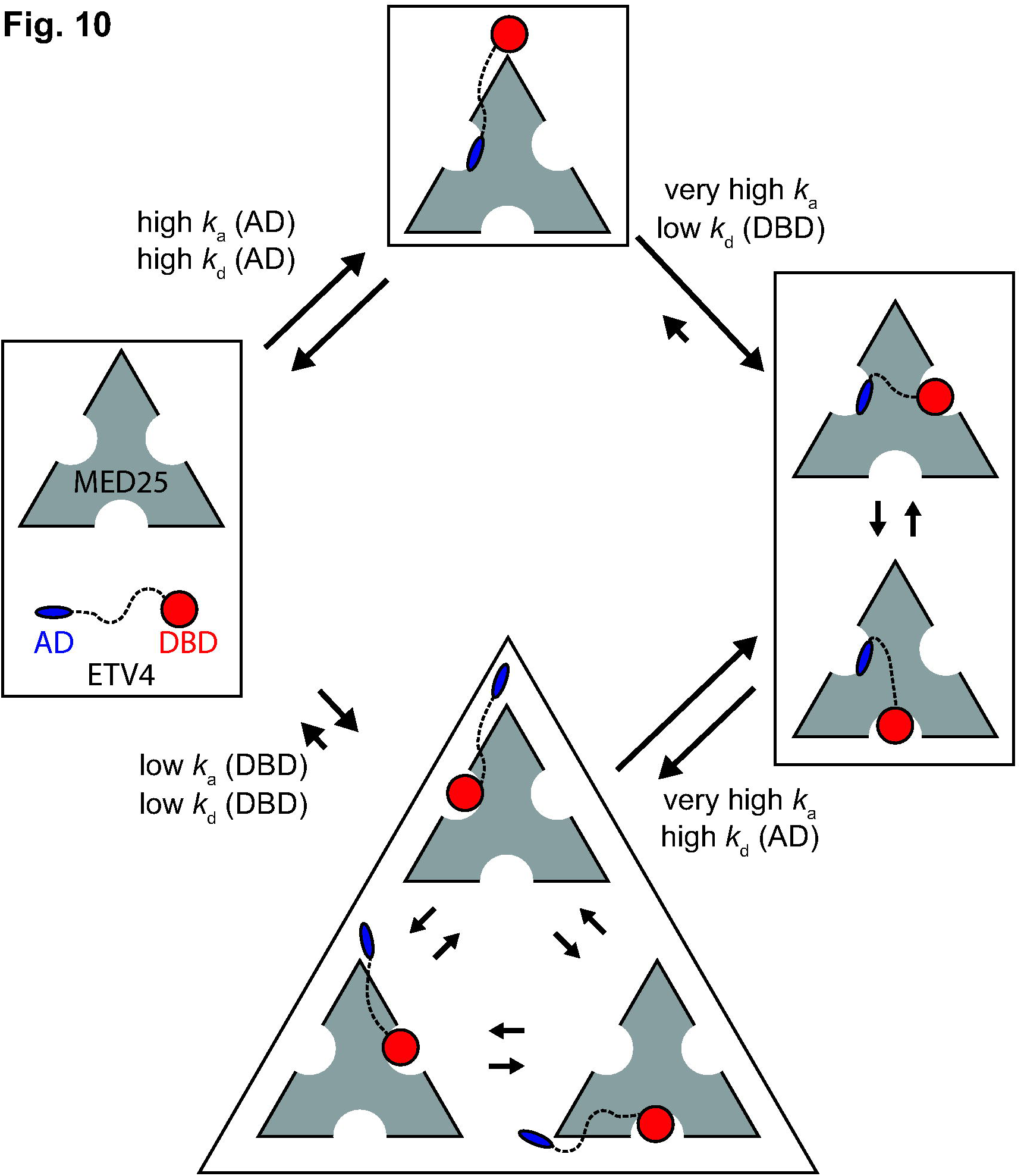
Model for the multivalent ETV4-MED25 interaction. In the predominant pathway (top) the first step of interaction involves the activation domain (AD) binding to MED25 Site 1 due to its drastically faster association compared to the interaction between MED25 and the DNA-binding domain (DBD, bottom). The association and dissociation rate constants, *k_a_* (AD), *k*_d_ (AD), *k_a_* (DBD), and *k*_d_ (DBD) for the first binding steps are assumed equivalent to those measured for the isolated AD-MED25 and DBD-MED25 interactions, respectively. In the second step of the predominant pathway, the DBD of the bound ETV4 can only bind to the two MED25 sites that are not occupied by the AD (right). The ETV4-MED25 interaction would then be in equilibrium between two states with the DBD alternately occupying Sites 2 and 3 on MED25. The predominant pathway for dissociation of ETV4-MED25 will be limited by the slower release of the DBD. ETV4 DBD binding to MED25 is not predicted to be the predominant first step for the interaction for full-length ETV4 and MED25 (bottom), due to the drastically slower association rate (bottom). However, in circumstances where the AD is bound to another protein, or in oncogenic translocations of ETV4 that do not have the N-terminal AD, binding of the DBD to MED25 would still be a viable mode of interaction. The size of the arrows indicating association and dissociation between the unbound and AD-MED25 or DBD-MED25 complexes, and dissociation between the ETV4-MED25 and AD-MED25 or DBD-MED25 complexes, is representative of the measured rate constants for these steps. Other arrows are qualitatively sized based on mutational data and interpretations, and should not be considered as measured values.

Interestingly, JUN/FOS factors also use a multivalent mechanism for interacting with MED25. FOS strongly interacts with MED25 by binding to Site 1, like ETV4 AD. JUN weakly interacts with MED25 by itself, but binds to MED25 Sites 2 and 3 when FOS is also present. Therefore, the multiple binding sites on MED25 provide a route for combinatorial interaction from a single transcription factor and may allow for cooperative recruitment of MED25 by multiple factors, an idea further explored below.

### Autoinhibition and regulation of ETV4

The conserved ETS domain results in similar DNA-binding properties for most ETS transcription factors [60]. However, evidence clearly indicates that individual ETS factors have distinct biological roles [61]; how is such specificity achieved? Part of this specificity can be ascribed to the diverse flanking inhibitory regions that are specific to individual subfamilies of ETS factors. In the ETS1 and ETV1/4/5 subfamilies, both inhibitory α-helices that pack against the ETS domain and intrinsically disordered sequences contribute to autoinhibition [54, 62]. The subfamily-specific α-helices also provide unique scaffolds for intermolecular interactions. In addition to MED25, USF1 relieves the DNA-binding autoinhibition of ETV4 [63], whereas, RUNX1 and PAX5 counter ETS1 autoinhibition [64, 65]. RUNX1 interacts with the ETS domain and flanking α-helices that are specific to ETS1 and ETS2 and blocks interaction between the disordered inhibitory region and the DNA-binding domain [66]. This DNA-binding activation in ETS1 is likely the reason that ETS1 specifically regulates ETS-RUNX composite DNA recognition sites in T-cells [67]. Here, we have demonstrated that MED25 interacts with the ETS domain and with H4, an inhibitory helix specific to the ETV1/4/5 subfamily. MED25 interaction with H4 relieves autoinhibition from H4 but does not appear to disturb inhibition from the disordered N-terminal inhibitory domain. Interestingly, acetylation of lysines relieves the inhibition from this disordered domain [54]. MED25 also interacts with CBP, a protein acetyltransferase [50]. Therefore, a hypothetical ETV4-MED25-CBP complex could maximally activate ETV4 DNA binding by relieving autoinhibition from both inhibitory domains and warrants further investigation. In conclusion, the ETV4-MED25 interaction is an example of how distinct flanking inhibitory sequences can set up specific protein partnerships that regulate the DNA-binding activity of a particular ETS factor.

### Multivalent AD-cofactor interactions

The multivalency of MED25 ACID with at least three distinct ETV4 binding sites could enable cooperative recruitment of MED25 to target genes. In PC3 cells, MED25 occupies genomic regions that were enriched for ETS and AP1 DNA-binding sequences. Thus, cooperative MED25 recruitment could be accomplished through interactions with AP1 and ETV4. We propose that FOS and ETV4 could simultaneously contact MED25 based on differences in binding modes; the strong interaction with FOS would occupy MED25 Site 1 while ETV4 DBD would bind to Sites 2 or 3 (Fig. S11). Many genomic regions have ETS and AP1 DNA-binding sites in close proximity that would allow for this concurrent interaction [57]. Such a combinatorial interaction could also explain the retained function of ETV1/4/5 truncations in prostate cancer that lack the N-terminal AD [31, 32]; the tighter binding FOS would outcompete ETV4 AD for interaction with MED25 Site 1, making the AD of ETV4 dispensable for the recruitment of MED25 to composite ETS-AP1 DNA sequences. The presence of multiple interaction sites on MED25 suggests that multiple transcription factors may cooperatively recruit MED25 to target genes.

In addition to the ETS factors and JUN/FOS described here, ATF6α, CBP, HNF4α, RARα, and SOX9 also interact with MED25 [43, 50, 51, 53]. ATF6α, CBP, and SOX9, bind to the ACID domain, though detailed structural and biochemical studies describing their binding sites have not been performed. Our genomic investigation did not suggest a broad role for these factors in recruiting MED25 in PC3 cells. However, given the plasticity of MED25 ACID binding sites used by ETV4, other ACID-binding factors may also be able to recruit MED25, along with ETV4, at select regions or in different cell types. The nuclear receptors (NRs), HNF4α and RARα, bind to MED25 through a NR box located outside of the ACID domain, which is thus distinct from Sites 1-3 described here. Therefore, cooperative recruitment of MED25 could be accomplished through simultaneous interaction with a nuclear receptor and any one of the ACID-interacting transcription factors. In particular, cooperative MED25 recruitment by ETV1/4/5 and androgen receptor (AR) could be an important mechanism for the oncogenic role of these transcription factors in prostate cancer; transcriptional coregulation by ETS and AR has been previously described [29, 68-70], although the potential importance of MED25 recruitment was not investigated.

MED25 ACID interacting with several different transcription factors is broadly illustrative of AD-cofactor interactions in general. Many cofactors are capable of interacting with ADs from multiple transcription factors. Conversely, most ADs can bind to multiple, unrelated cofactors. With this plasticity of binding options, what is the molecular basis for high affinity interactions that are specific to a single AD-cofactor partnership? Several examples, including our findings with ETV4, suggest that multivalency is one route for forming specific, higher-affinity AD-cofactor interactions. For example, two ADs from p53 form bivalent interactions with both the TAZ1 and TAZ2 domains of CBP by binding to opposite faces of these domains [21]. The p53 ADs also interact with at least two other CBP domains [24], so the tetrameric form of p53 has been predicted to form a highly multivalent interaction with full-length CBP. RAD4, RAD34, and TFIIE utilize bivalency to strongly bind to the PHD domain of the general transcription factor TFIIH [22, 23]. The GCN4-MED11 interaction may be most reminiscent of MED25 and ETV4; two GCN4 ADs interact with three different activator-binding domains in MED11 [20].

Thus, ETV4-MED25 interactions described here extend the emerging picture of multivalent interactions between transcription factors and transcriptional coactivators in the assemblage of transcriptional machinery. Multiple interacting surfaces enables the generation of higher-affinity and specific interactions while only a single binding site mediates lower affinity interactions; the avidity between the transcription factor and the cofactor can be altered by modulating the number of potential interacting surfaces on each protein. In the case of oncogenic proteins, such as ETV4 as well as related factors ETV1 and ETV5, this understanding of multivalent interactions may enable a multipronged approach for therapeutic strategies.

## Materials and Methods

### Protein Expression & Purification

The genes encoding full-length ETS factors and truncated ETV4 fragments were cloned into the pET28 (Novagen) bacterial expression vector using sequence and ligation independent cloning (SLIC) [71]. ETV4^337-436^ that was used as a ligand in biolayer interferometry was cloned into a custom-made vector, described below. Plasmid of human MED25 cDNA was ordered from GE Dharmacon. The gene encoding MED25 ACID (residues 391-553) was cloned into pET28 to enable protein production for NMR spectroscopy, and into a custom-made vector to express ligand protein for biolayer interferometry. This vector was based on a pGEX backbone with N-terminal GST, Avitag, and HIS_6_ tags separated by thrombin and TEV cleavage sites, and followed by a Precission protease site before MED25 ACID. Plasmids of JUN and FOS in the pET28 bacterial expression vector were a kind gift from Dr. Peter Hollenhorst.

All proteins, except JUN and FOS, were expressed in *Escherichia coli* BL21 (λDE3) cells. MED25 ACID, ETV4^1-164^, ETV4^43-84^, and ETV4^337-436^ expressed into the soluble fraction of cells, and were grown in 1L cultures of Luria broth (LB) at 37**°** C to OD_600_ ∼ 0.7 – 0.9, induced with 1 mM isopropyl-β-D-thiogalactopyranoside (IPTG; Gold Biotechnology), and grown at 30**°** C for ∼ 2-3 hrs. For MED25 ACID or ETV4^337-436^ protein used as the ligand in biolayer interferometry, 1 mL of 50 mM D-biotin (Fisher Scientific) was added at the induction point. Cells were centrifuged at 6,000 rpm in a JLA 8.1 rotor (Beckmann), resuspended in a buffer containing 25 mM Tris, pH 7.9, 200mM NaCl, 0.1 mM ethylenediaminetetraacetic acid (EDTA), 2 mM 2-mercaptoethanol (βME), and 1 mM phenylmethanesulfonylfluoride (PMSF), and snap-frozen in liquid nitrogen. After 3 to 5 freeze-thaw cycles, the cells were lysed by sonication and ultracentrifuged for 30 min at 40k rpm in a Beckman Ti45 rotor. The soluble fraction was then loaded onto a Ni^2^+ affinity column (GE Biosciences) and eluted over 20 column volumes of a 5-500 mM imidazole (Sigma) gradient. For MED25 ACID used in biolayer interferometry, protein eluted from the Ni^2^+ column was loaded onto a GST-affinity column (GE) and eluted with the same buffer with 15 mM glutathione (Sigma) added. Elutions were then dialyzed overnight into 25 mM Tris pH 7.9, 10% glycerol (v:v), 1 mM EDTA, 50 mM KCl and 1 mM dithiothreitol (DTT; Sigma). After ultracentrifugation for 30 min at 40k rpm in a Beckman Ti45 rotor, proteins were purified over a SP-sepharose cation exchange column (GE) (ETV4^337-436^ and MED25 ACID) or a Q-sepharose anion exchange column (GE) (ETV4^1-164^ and ETV4^43-84^) using a 50-1000 mM KCl gradient. Proteins were then further purified over a Superdex 75 gel filtration column (GE) in 25 mM Tris pH 7.9, 10% glycerol (v:v), 1 mM EDTA, 300 mM KCl, and 1 mM DTT. Purified proteins were concentrated with a 3, 10, or 30 kDa molecular weight cutoff (MWCO) centricon device (Sartorius), snap-frozen in liquid nitrogen and stored in single-use aliquots at -80 °C for subsequent NMR, EMSA, or biolayer interferometry studies.

Full-length ETV4 (ETV4^1-484^) and ETV4^165-484^ expressed into the insoluble fraction and were readily induced by autoinduction [72]. Briefly, bacteria in 250 mL of autoinduction media were grown in 4 L flasks at 37° C to an OD600 ∼ 0.6 – 1. The temperature was then reduced to 30° C and cultures were grown for another ∼ 12 – 24 hr. Final OD_600_ values were typically ∼ 6 – 12, indicating robust autoinduction. Harvested cells were resuspended as described above, sonicated and centrifuged at 15k rpm in a JA-17 rotor (Beckman) for 15 min at 4° C. The soluble fraction was discarded and this procedure was repeated with the pellet / insoluble fraction twice more to rinse the inclusion bodies. The final insoluble fraction was resuspended with 25 mM Tris pH 7.9, 1 M NaCl, 0.1 mM EDTA, 5 mM imidazole, 2 mM βΜΕ, 1 mM PMSF, and 6 M urea. After sonication and incubation for ∼ 1 hr at 4**°** C, the sample was centrifuged for 30 min at 40k rpm and 4**°** C. The soluble fraction was loaded onto a Ni^2^+ affinity column and refolded by immediately switching to the same buffer except without urea. After elution with 5-500 mM imidazole, the remaining purification steps (anion-exchange and size-exclusion chromatography) were performed as described above.

Full-length JUN and FOS proteins expressed into the insoluble fraction using IPTG induction and were expressed and purified as for insoluble ETV4 proteins, described above, with the following exceptions. FOS was expressed in Rosetta 2 cells for supplementation of rare Arg tRNAs. Inclusion bodies were purified and solubilized as described above, then JUN and FOS were combined for JUN/FOS heterodimers (or kept separate for JUN homodimers or FOS monomers), diluted to 200 ng/μL (total protein), then dialyzed for at least 3 hr each against the following three buffers (in sequential order): (1) 25 mM Tris pH 6.7, 0.1 mM EDTA, 10% glycerol, 5 mM BME, 1 M NaCl, 1M Urea; (2) same as (1) but without urea; (3) same as (2) but with NaCl reduced to 100 mM. Refolded samples were then purified using Ni^2^+ affinity column and gel-filtration column as described above.

### Biolayer Interferometry

Data were collected with an Octet Red96 instrument (ForteBio) and processed with Octet Data Analysis software (v8.1, ForteBio). MED25 ACID (residues 391-553) was used as the ligand for all interactions except for with ETV4^337-436^ as this truncation of ETV4 displayed too much non-specific interaction with the biosensor tip to be used as the analyte. Biotinylated ligand (500 nM) was immobilized using high precision streptavidin sensors (ForteBio). Protein concentrations were determined after thawing each aliquot of protein, using the Protein Assay Dye Reagent (Bio-Rad). Interaction experiments were conducted using 25 mM Tris pH 7.9, 150 mM NaCl, 10 mM DTT, 5 μg/mL bovine serum albumin (Sigma) and 0.05% (v:v) Tween20 (Sigma). Biosensors were dipped in various concentrations of the analyte of interest to measure association, and transferred back to buffer wells for monitoring dissociation.

For a more quantitative analysis, six concentrations of analyte, with exact concentrations varying dependent on the affinity of the specific analyte-ligand combination, were fit using a global (full) analysis (simultaneous, constrained fit of all six sensorgrams). The reported kinetic rate constants (*k_a_* and *k*_d_) were determined from the fit of a 1:1 binding model for ETV4^43-84^ and ETV4^1-164^, and a 2:1 (ETV4:MED25) binding model for all full-length ETS factors, ETV4^165-484^, and ETV^337-436^. The decision to use either of these binding models was determined empirically based on chi-squared values, as evidence from NMR spectroscopy and protein-docking experiments also suggested that ETV4^337-436^ interacts with MED25 using a multivalent mode. The equilibrium dissociation constant (*K_D_*) was derived from the kinetic constants. Use of either the 1:1 or 2:1 binding models resulted in relatively similar *K*_D_ values for all interactions tested. For example, the full-length ETV4-MED25 interaction results in a *K*_D_ of 5 ± 1 nM using the 1:1 binding model and *K*_D_ values of 7 ± 3 and 28 ± 7 using the 2:1 binding models. Agreement was similar for other ETV4 fragments and ETS factors. Therefore, the use of either model supports the conclusions that ETV1/4/5 subfamily factors bind MED25 more tightly than other ETS factors and that full-length ETV4 binds more tightly to MED25 than the AD or DBD alone. All constants were averaged separately from replicate experiments, therefore the reported *K*_D_ value may not exactly equal *k*_d_ / *k*_a_. Mean and standard deviation of *K*_D_, *k*_a_, and *k*_d_ values from at least three independent experimental replicates are displayed in figures and tables.

### NMR Spectroscopy

^1^H-^15^N HSQC measurements were recorded on a 500MHz Varian Inova spectrometer at 25**°** C with protein in the following sample buffer: 20mM sodium phosphate, pH 6.5, 200 mM NaCl, 2 mM DTT, 1 mM EDTA, and 10% D_2_O. Proteins were either purified into NMR sample buffer using a size-exclusion column, or buffer-exchanged using centricon spin columns. Labeled protein was concentrated to ∼ 0.2 – 0.25 mM and titrated with 0.2, 0.5, and 1.2 molar equivalents of unlabeled protein. Upon addition of each titration equivalent, the entire sample was removed and concentrated back down to approximately the same starting volume. Assignments were transferred from previous work for MED25 ACID [9, 73] and ETV4 337-436 [54]. NMR data were processed and analyzed using Vnmr (Varian) and Sparky (UCSF) [74]. Chemical shift perturbations (Δδ = [(Δδ_H_)^2^ + (0.2Δδ_N_)^2^]½) and relative peak intensities for ETV4 and MED25, respectively, that were above the median and in the top ten percent for chemical shift perturbations, or below the median and in the lowest ten percent for relative peak intensities, were colored onto the structures of ETV4 (PDB: 4UUV)[56] or MED25 (PDB: 2KY6)[9] using Pymol (v1.7.0.5 enhanced for Mac, Schrödinger).

### Electrophoretic Mobility Shift Assays (EMSAs)

DNA-binding assays of ETS factors utilized a duplexed 27-bp oligonucleotide with a consensus ETS binding site: 5’-TCGACGGCCAAGCC**GGAA**GTGAGTGCC-3’ and 5’-TCGAGGCACTCACTTCCGGCTTGGCCG-3’. Boldface **GGAA** indicates the consensus ETS binding site motif. Each of these oligonucleotides, at 2 μM as measured by absorbance at 260 nM on a NanoDrop 1000 (Thermo Scientific), were labeled with [γ-^32^P] ATP using T4 polynucleotide kinase at 37 °C for 30 minutes. After purification over a Bio-Spin 6 chromatography column (Bio-Rad), the oligonucleotides were incubated at 100 °C for 5 minutes, and then cooled to room temperature over 2 hours. The DNA for EMSAs was diluted to 1 × 10^-12^ M and held constant. ETV4 and EHF concentrations ranged was diluted to 5 × 10^-7^ and held constant while MED25 was diluted from 5 × 10^-8^ to ∼ 1 × 10^-9^ M. For measuring ETS-DNA *K*_D_ values, MED25 was held constant at 5 × 10^-7^ and ETS factors were diluted from 1 × 10^-7^ to ∼ 1 × 10^-11^ M. Protein concentrations were determined after thawing each aliquot of protein, using the Protein Assay Dye Reagent (Bio-Rad). The binding reactions were incubated for 45 minutes at room temperature in a buffer containing 25 mM Tris pH 7.9, 0.1 mM EDTA, 60 mM KCl, 6 mM MgCl_2_, 200 □g/mL BSA, 10 mM DTT, 100 ng/□L poly(dIdC), and 10% (v:v) glycerol, and then resolved on an 8% (w:v) native polyacrylamide gel at room temperature. The ^32^P-labeled DNA was quantified on dried gels by phosphorimaging on a Typhoon Trio Variable Mode Imager (Amersham Biosciences). Equilibrium dissociation constants (*K*_D_) were determined by nonlinear least squares fitting of the total protein concentration [P]_t_ versus the fraction of DNA bound ([PD]/[D]_t_) to the equation [PD]/[D]_t_ = 1/[1 + *K*_D_/[P]_t_)] using Kaleidagraph (v. 3.51; Synergy Software). Due to the low concentration of total DNA, [D]_t_, in all reactions, the total protein concentration is a valid approximation of the free, unbound protein concentration. Reported **K*_D_* values represent the mean and the standard deviation from at least four independent experiments.

### Protein Docking

Modeling for ETV4 AD- and DBD-MED25 ACID interactions were performed using the ZDOCK server (http://zdock.umassmed.edu) [55]. PDB files 4UUV [56] and 2KY6 [9] were used as inputs for ETV4 DBD and MED25 ACID, respectively. An α-helix with the sequence “LSHFQETWLAEA” was generated using Pymol and used as the input for ETV4 AD based on the conserved sequence in ETV5 becoming more helical when interacting with MED25 ACID [7]. From our mutational data, we selected ETV4 residues Phe60 and Trp64, and Phe428 and Trp432 to be involved in the interactions between AD-MED25 ACID and DBD-MED25 ACID, respectively. We did not select any ETV4 residues to be blocked from the interface. Our NMR data showed broad perturbations of residues on multiple faces of MED25 ACID by both ETV4 AD and DBD; therefore, we did not select any MED25 residues to be involved, nor to be blocked, in these predicted interactions.

### Statistical Analysis

An unpaired Mann-Whitney test was used to calculate *p* values using Graph Pad Prism (v.6). Values less than 0.05 were considered significant and are indicated by “*”; whereas, values greater than 0.05 were not considered significant and are indicated by brackets without an asterisk. Replicate numbers are indicated by the number of dots in each bar graph, and are included in Tables S1, S2, and S6.

### Cell culture and viral expression

PC3 cells were obtained from American Type Culture Collection and cultured accordingly. Full-length MED25 cDNA was cloned into pQCXIH (Clontech) with an added C-terminal 3x FLAG tag. Oligonucleotide sequences for MED25 shRNA hairpin design are as follows, with targeting sequences in lower case: MED25a_fwd-CCGGtgattgagggtacggccaaTTCAAGAGAttggccgtaccctcaatcacaTTTTTG MED25a_rev-AATTCAAAAAtgtgattgagggtacggccaaTCTCTTGAAttggccgtaccctcaatca MED25b_fwd-CCGGtcaaaggcctctaccgcatTTCAAGAGAatgcggtagaggcctttgagaTTTTTG MED25b_rev-AATTCAAAAAtctcaaaggcctctaccgcatTCTCTTGAAatgcggtagaggcctttga MED25 shRNA hairpins were cloned into the pLKO.1 lentiviral expression vector [75], and expression and infection of lentivirus performed following standard protocol. ETV4 shRNA retroviral constructs were previously described, and retrovirus production and infections were carried out following previously published methodology [76]. Whole-cell extracts from cells expressing control shRNA constructs, ETV4 shRNA or MED25 shRNA were run on SDS-PAGE gels and blotted to nitrocellulose membranes following standard procedures. Antibodies used for immunodetection were ETV4, ARP32262 (Aviva Systems Biology); MED25, ARP50699_P050 (Aviva Systems Biology); and beta-Tubulin, sc-55529 (Santa Cruz).

### Chromatin Immunoprecipitation (ChIP) and ChIPseq analysis

ChIPs were performed as described previously [67], with the following modifications. Cross-linked chromatin was sheared with a Branson sonifier and magnetic beads were washed with buffer containing 500 mM LiCl. Antibodies used for ChIP were: MED25, anti-Flag M2, (Sigma Life Sciences); ETV4, ARP32262 (Aviva Systems Biology). ChIPseq libraries were prepared using the NEBNext^®^ ChIP-Seq Library Prep Master Mix Set for Illumina (NEB, E6240) and run on a Hiseq2000 sequencer. Two ETV4 and two MED25 ChIPseq libraries were generated from biological replicates and analyzed as replicates with one input control library. Sequence reads were aligned with Bowtie [77] to human genome HG19 and enriched regions (peaks) determined using the Useq analysis package [78], at an FDR of 1% and Log2ratio of 2 with input DNA library used as control sample and either ETV4 or MED25 aligned reads as treatment sample. Shared regions between ETV4 and MED25 enriched regions were defined using IntersectRegion in Useq package with no gap. Heatmaps of shared regions were generated with DeepTools [79] using a bedfile corresponding to coordinates from the MED25-ETV4 shared regions from MED25 dataset, and bigwig files generated from the aligned ETV4, MED25 and input ChIPseq reads for one replicate. Data were aligned using the center point of this shared peak bedfile for all three maps. Enriched regions were also visualized graphically using IGV [80].

Overrepresented DNA sequences present in the ETV4 and MED25 enriched regions were determined using the MEME-ChIP program [81] (http://meme-suite.org) using default settings except for following parameters for MEME-1) any number of repetitions for site distribution; and 2) maximum site width of 13. The central 100 bp of the ETV4 and MED25 enriched regions were interrogated by MEME-ChIP to find centrally located binding sites within the regions. We used these tighter peak coordinates with IntersectRegions to generate a more stringent set of shared ETV4-MED25 regions resulting in 611 shared regions to interrogate with MEME-ChIP. A set of size-matched random regions for comparison to the 611 ETV4 and MED25 shared regions was generated using the shuffle command in bedtools (http://bedtools.readthedocs.io/en/latest/). The 611 shared regions and 611 randomly generated regions were searched for the occurrences of C(A/C)GGAA using the FIMO program from MEME suite [82].

### RNAseq sample preparation and analysis

RNA samples for each shRNA construct (ETV4a, ETV4b, MED25a, MED25b, lenti-luciferase shRNA) were analyzed as biological triplicates except for ETV4 shRNA retroviral control, which had two biological replicates. RNA was prepared from cells at day 7 post-infection using the Qiagen RNAeasy Mini Kit. Samples were treated on column with DNase to remove genomic DNA. Samples were depleted of ribosomal DNA using a bead-based method (Illumina TruSeq Stranded RNA Kit with Ribo-Zero Gold) prior to library construction. Library construction was performed using the Illumina TruSeq Stranded RNA Kit and sequenced using the Illumina Hiseq2000 sequencer. Sequence reads were aligned to hg19 using novoalign (Novocraft) and differentially expressed transcripts determined with drds analysis package in Useq. Expression changes of an absolute log 2 fold change > 0.585 and FDR <1%, relative to control shRNA, were counted as significant. In order to be considered affected by both ETV4 and MED25 shRNA, the expression of a gene had to be significantly affected by all four shRNA constructs (ETV4a, ETV4b, MED25a, MED25b).

### qPCR analysis of ChIP DNA and RNA

Primer-BLAST [83] was used to generate primer sets for amplification of enriched regions; primer sequences and their coordinates are provided in supplemental table S7, and regions are also denoted as red bars in IGB tracks (Fig. 8b and Fig. S8b). qPCR of ChIP DNA was performed using Roche FastStart Essential DNA Green Master and run on a Roche Lightcycler 96. Serially diluted input was used to create a standard curve for absolute quantitation of amplified regions from ChIP DNA. PCRs for each sample and primer pair were run as triplicates and signal averaged over the three values. Data are displayed in graphical form as a ratio of the signal of the target region over the signal of a negative control genomic region. An input sample was also subject to same qPCR reactions graphed to confirm validity of negative control region. For all primer pairs, the input enrichment value was approximately one.

For qPCR analysis of RNA samples, total cDNA was generated from RNA using iScript Reverse Transcription Supermix (Bio-Rad) and used as a DNA template for qPCR with Roche FastStart Essential DNA Green Master. Primers pairs for each gene assayed were designed with Primer-BLAST against the corresponding mRNA. Relative gene expression was calculated using the ΔΔCq method, with GAPDH as an internal reference. Reactions were performed in triplicate for each sample and ΔΔCq values averaged for each knockdown RNA sample relative to the average ΔCq value for the control with standard deviation. Expression levels of individual genes in samples from ETV4 and MED25 shRNA knockdown samples are reported as values relative to control shRNA.

## Acknowledgments

We thank Peter Hollenhorst for providing *JUN* and *FOS* plasmids and for critical review of the manuscript. This study was funded by the National Institutes of Health (R01GM38663 to B.J.G.), the Canadian Cancer Society Research Institute (CCSRI 2011-700772 to L.P.M.), and the Canadian Institutes of Health Research (CIHR MOP-136834 to L.P.M.). Instrument support was provided by CIHR, the Canada Foundation for Innovation (CFI), the British Columbia Knowledge Development Fund (BCKDF), the UBC Blusson Fund, and the Michael Smith Foundation for Health Research (MSFHR). Support to B.J.G. from the Huntsman Cancer Institute/Huntsman Cancer Foundation and the Howard Hughes Medical Institute is acknowledged. Shared resources at the University of Utah, including the NMR facility, were supported by the National Institutes of Health (P30CA042014 to the Huntsman Cancer Institute).

## Abbreviations Used

AD: activation domain
ACID: activator-interacting domain
ChIP-Seq: chromatin immunoprecipitation followed by deep sequencing
DBD: DNA-binding domain
ETS: E26-transformation specific
MED25: Mediator subunit 25
HSQC: heteronuclear single quantum coherence
AP1: activator protein 1
AR: androgen receptor

